# Noradrenergic Regulation of Two-Armed Bandit Performance

**DOI:** 10.1101/2020.11.13.382069

**Authors:** Kyra Swanson, Bruno B. Averbeck, Mark Laubach

**Author notes:** Correspondence concerning this article should be addressed to Mark Laubach, Department of Neuroscience, American University, 4400 Massachusetts Ave NW, Washington, DC 20016.

## Abstract

Reversal learning depends on cognitive flexibility. Many reversal learning studies assess cognitive flexibility based on the number of reversals that occur over a test session. Reversals occur when an option is repeatedly chosen, e.g. eight times in a row. This design feature encourages win-stay behavior and thus makes it difficult to understand how win-stay decisions influence reversal performance. We used an alternative design, reversals over blocks of trials independent of performance, to study how perturbations of the medial orbital cortex and the noradrenergic system influence reversal learning. We found that choice accuracy varies independently of win-stay behavior and the noradrenergic system controls sensitivity to positive feedback during reversal learning.

---

Reversal learning tasks are one of three types of behavioral tasks used to study behavioral flexibility, along with attentional shifting and rule switching (Tait et al., 2018). Reversal learning requires participants to remap associations between either stimuli or actions and their outcomes. A number of disorders have been reported to negatively influence reversal learning (Peterson et al., 2009; Reddy et al., 2016; Remijnse et al., 2006; Waltz & Gold, 2007), and the task has been suggested as a core method for preclinical evaluations of drugs used for treating psychiatric disorders (Powell & Ragozzino, 2017).

Some researchers have described each decision in a reversal learning task in terms of Win-Stay/Lose-Shift (WSLS) strategies (Bari et al., 2010; Dalton et al., 2014, 2016; Jang et al., 2015). The Win-Stay (WS) strategy exploits a previously-rewarded choice, while the Lose-Shift (LS) strategy involves exploration of a different option after reward omission (Estes, 1950). Probabilistic outcomes that force animals to integrate multiple previous trials to guide choice increase the difficulty of the task, and the task design has commonly been refered to as “probabilistic reversal learning”. Other studies have thus referred to task design as a “bandit” task--a reinforcement learning paradigm. Within this framework, the focus is on changes in two variables: learning rate, which describes how quickly the subject incorporate new evidence into its decisions, and inverse temperature, which describes the subject’s confidence in the decision (Groman et al., 2016; Metha et al., 2019; Sutton & Barto, 2018). These two approaches, WSLS and reinforcement learning, are rarely combined to understand behavioral mechanisms or neuronal measures of reversal learning (Worthy & Maddox, 2014). One recent attempt at doing this was reported by Harris et al. (2020), who found evidence for a dissociation among measures of accuracy, latency, WSLS, and reinforcement learning during the initial discrimination learning and reversal learning of a visually guided probabilistic learning task.

Several neurotransmitter systems (Robbins & Roberts, 2007) and selective regions of frontal cortex (Izquierdo et al., 2017) have been implicated in behavioral flexibility, and bandit performance in particular. Among these systems, the role of the norepinephrine system and its actions in the orbitofrontal cortex are unclear. Neural recording studies have established that the orbitofrontal cortex (OFC) is important for maintaining predictive stimulus-outcome associations (e.g. Schoenbaum et al., 1999). Furthermore, OFC lesions or reversible inactivations have consistently impaired stimulus-outcome remapping in reversal learning tasks (e.g. Chudasama & Robbins, 2003). This part of the frontal cortex is therefore an important target for understanding how norepinephrine modulates neural processing during two-armed bandit performance.

The noradrenergic system also has an established role in reversal learning (Seu et al., 2009). Tonic norepinephrine (NE) activity in the OFC has been proposed to modulate cognitive flexibility by allowing the formation of novel contexts and associations (Sadacca et al., 2017; Wilson et al., 2014). The α2 antagonist yohimbine reduces availability of NE receptors throughout the brain and may lead to reduced persistent activity in the frontal cortex (Kovács & Hernádi, 2003; Zhang et al., 2013).. Yohimbine may therefore impair the ability to form new spatial/action-outcome contingencies (Sadacca et al., 2017; Wilson et al., 2014), and could alter how animals learn from feedback (Jepma et al., 2016). No published study has examined the impact of yohimbine on performance in a two-armed bandit task or how NE-selective drugs alter WSLS strategies or models of reinforcement learning.

In the present study, we evaluated how rats perform a version of the two-armed bandit task that has blocked option-outcome reversals (Costa et al., 2015). This allowed us to investigate the relationships between WSLS strategy, reinforcement learning, and flexibility, and also to examine within-session changes under a range of outcome probabilities. We further examined the role of the medial OFC using reversible inactivation methods and evaluated effects of systemic and intra-cortical antagonism of NE on two-armed bandit performance. There were three main findings from our study. First, rats maintained their WSLS strategy across different reward probabilities, but adapted their strategy with experience in each session. Second, reversible inactivation of the mOFC decreased the animals’ accuracy in performing the task and their sensitivity to negative feedback. Third, NE antagonism, systemically but not intracranially in mOFC, decreased performance accuracy, reduced sensitivity to positive feedback, and decreased the inverse temperature parameter from the reinforcement learning algorithm.

## Methods

Procedures were approved by the Animal Use and Care Committee at American University and conformed to the standards of the National Institutes of Health as outlined in the “Guide for the Care and Use of Laboratory Animals” published by the Public Health Service.

### Subjects

Twenty four male Long-Evans rats (300-350g) were obtained from the NIH animal colony, Charles River, or Envigo. Animals were housed individually and kept on a 12/12 h light/dark cycle. Animals had regulated access to food to maintain their body weights at approximately 90% of their free-access weights. Seven of these animals were unable to reach the initial accuracy criterion in the spatially-guided deterministic blocked bandit design (described below) and were removed from the study. Seventeen rats in total were tested in the uncertainty experiment. From this group, 11 rats were tested with 2mg/kg systemic yohimbine. Five of those 11 were surgically implanted with cannulae for intra-cortical infusions of muscimol. An additional 4 Long-Evans rats (Charles River) were trained and tested with yohimbine and muscimol but did not participate in the uncertainty experiment.

### Behavioral Apparatus

All animals were trained in sound-attenuating behavioral boxes (ENV-018MD-EMS: Med Associates). A light-pipe lickometer was located on one side of the box, and contained a 5/16” sipper tube recessed behind a photobeam (ENV-251M: Med Associates). The tip of the sipper tube was 6.5 cm from the floor of the box. A green LED light (4 cm) was placed above the spout. The opposite wall had two nosepoke ports aligned horizontally 4.5 cm from the floor, 12.5 cm apart, with IR beam break sensors on the external side of the wall. Behavioral devices were controlled and data was collected using custom-written code for the MedPC system, version IV (Med Associates). Visual stimuli were presented using custom-made LED matrices (Swanson et al., 2021). Visual stimulus devices were placed above each nosepoke port, outside the box, to signal active trials. Positive reward feedback was presented through either a 1-sec 4.5 kHz SonAlert tone (Mallory SC628HPR) or a mechanical relay which clicked three times in quick succession. Negative omission feedback was presented through with a 1-sec 2.9 kHz SonAlert tone (Mallory SC105R). The devices were placed in opposite corners of the box behind the spout.

### Training Procedure

Rats were first given access to 25 ml of 16% sucrose in their home cage once a day over a two day period. They were then acclimated to the operant chamber with unrestricted access to liquid sucrose from the lickometer in 30 minutes behavioral sessions. Fluid presentation was indicated by activation of the fluid pump, the relay auditory stimulus, and illumination of a 5 mm green LED located above the lickometer. Once the rats licked more than 1000 times in each of two consecutive behavioral sessions, the rats were trained to nosepoke at either the left or right ports in a counterbalanced manner. The visual stimulus above the nosepoke was fully illuminated at the start of each trial. The opposite nosepoke was inaccessible. After 2-5 training sessions, in which the open nosepoke alternated each day, we then opened both nosepokes at once while retaining the visual cues. The rats had to respond at the visually-cued nosepoke for 2-4 days. If the unilluminated port was chosen, the animals would be presented with the 2.9 kHz tone and would have to lick at the spout to begin a new trial.

After rats responded selectively to the illuminated port, we introduced 30-trial blocks with fixed locations of the cued and rewarded ports. After every 30 trials, the cued and rewarded ports switched to the alternate side. On the very first trial in the session, both ports were illuminated, and the option chosen on this trial became the “correct”, illuminated side for the remainder of the block. Finally, the task transitioned from visually and spatially guided to purely spatially guided behavior. The difference in the number of illuminated LEDs in the LED matrices (Swanson et al., 2021) between the correct and incorrect side was slowly decreased over several sessions, starting from 10 or 8 versus 0 LEDs, until both cues had an equal number of LEDs illuminated for each trial.

Once rats were able to perform the spatially-guided deterministic blocked bandit at an accuracy of approximately 80%, the rats were further tested for 3 days, and then tested for three days each on 90/10, 80/20, and 70/30 reward schedules in order. We did not test them on a 60/40 reward schedule because their accuracy for the 70/30 test sessions was at or below the 65%, and thus near chance.

### Systemic Drug Injections

Following initial testing under different reward schedules, rats were either returned to a deterministic schedule for 1-2 days before being challenged with yohimbine or implanted with infusion cannulas (as described below). Rats that received drug cannulas were tested first with central infusions and then with systemic injections of yohimbine. For systemic drug testing, rats were first injected with a physiological saline volume control, and the next day were tested with 2 mg/kg yohimbine (Yobine: Akorn Pharmaceuticals). Both treatments were administered intraperitoneally under isoflurane 20 minutes before testing. There was no difference in behavior between a gas-only pretest and the saline session.

### Surgery

Nine rats were surgically implanted with bilateral cannulae. Anesthesia was induced by an injection of diazepam (5 mg/kg, IP) and maintained with isoflurane (4.0%; flow rate 4.5 cc/min). Standard stereotaxic methods were used to implant 26-gauge guide cannulae (PlasticsOne) bilaterally, targeting the medial orbitofrontal cortex (as in Swanson et al., 2019). Depth was calculated from the brain surface using a posterior angle of 12° and a lateral angle of 30° (AP: 3.6mm ML: 1.2mm DV: 2.0mm). Infusions were made using 33 gauge cannula (PlasticsOne), which extended 0.5 mm past the end of the guide cannulae. We used the term medial orbitofrontal cortex (mOFC) to refer to the general region of the frontal cortex examined in this study, as in a recent study from our group (Swanson et al., 2019). This term was used because all infusion cannula were localized in the medial frontal cortex anterior to the rhinal sulcus. We note that mOFC is not a homogeneous region, and includes the medial orbital cortex (ventral), rostral prelimbic cortex (dorsal), and medial portion of ventral orbital region (lateral).

Rats were allowed 7 days of recovery before returning to behavioral testing. Intra-cortical drug infusions were done under isoflurane anesthesia as in previous studies from our lab (e.g. Swanson et al., 2019). Rats were acclimated to testing after 10 minutes under gas anesthesia prior to testing. Rats showed no differences in behavioral performance (trials performed, accuracy) after 1-2 “gas control” sessions. For muscimol testing, rats were infused with 0.5 μL of muscimol (Tocris) or physiological saline with a flow rate of 0.25 μL/min. Seven of the animals were tested with 0.05 μg/μL muscimol. Two did not perform the task at this dosage, so instead we report their behavior under 0.01 μg/μL. Four of the rats were also infused with 2 μL of either physiological saline or 5 μg/μL yohimbine (Tocris) with a flow rate of 0.5 μL/min as previously reported (Caetano et al., 2012). Animals were tested 15 minutes after infusion. For both muscimol and yohimbine tests, infusion cannula were left in place for 2 minutes after the infusions to allow for diffusion.

### Confirmation of Cannula Placement

After completion of the experiment, animals were anesthetized with isoflurane gas and injected intraperitoneally with Euthasol (100 mg/kg). Animals were transcardially perfused with 200 ml of chilled (4°C) physiological saline solution followed by 200 ml of chilled (4°C) 4% paraformaldehyde. Brains were removed and cryoprotected using a solution containing 20% sucrose, and 20% glycerol. Brains were then cut into 50 μm-thick coronal slices using a freezing microtome (Hacker). Brain sections were mounted onto gelatin-coated slides and Nissl stained via treatment with 0.05% thionin. Thionin-treated slices were dehydrated through a series of alcohol steps, then covered with Clearium and coverslipped. Sections were imaged using a Tritech Research scope (BX-51-F), Moticam Pro 282B camera, and Motic Images Plus 2.0 software. The most ventral point of the injection tract was compared against (Paxinos & Watson, 2007) to estimate brain atlas coordinates.

### Data Analysis

Behavioral data were saved in standard MedPC data files and were analyzed using custom-written code in the Python (Python Software Foundation, https://www.python.org/) and R (The R Project, https://www.r-project.org/) languages, maintained using Anaconda (https://www.anaconda.com). Analyses were conducted using Jupyter notebooks (https://jupyter.org/). The relationships between WSLS strategy, accuracy, and reward uncertainty were evaluated using analysis of variance (ANOVA) and linear regression. Within-session effects were evaluated using ANOVA, and block analysis was done using repeated-measures ANOVA. Post hoc testing used either Tukey’s Honest Significant Difference (HSD) test for standard ANOVA or paired t tests within a given level of block for repeated-measures ANOVA. Statistical testing and post hoc analysis were done using the default functions in R, e.g. aov.

Accuracy was defined as the percentage of trials directed toward the option with the higher likelihood of dispensing reward. Win-Stay values were determined by calculating the proportion of stay trials following every win. The inverse of this proportion is the likelihood of demonstrating Win-Shift behavior. Likewise, Lose-Shift values were determined by calculating the proportion of shift trials following every omitted reward. The inverse of the proportion is the likelihood of demonstrating Lose-Stay behavior. For ease of computation, the first trial in every session was considered a “Stay” trial in reference to a hypothetical 0th trial.

Perseverative errors were defined as responses to the previously high-value option immediately after a reversal, before the first response to the new high-value option (Caetano et al., 2012). Please note that this measure includes rewarded errors given the probabilistic nature of the task.

Change points for choices during each block were calculated using the cpt.mean function in the *changepoint* package for R (Killick and Eckley, 2014). Change points were estimated based on the likelihood ratio test statistic. The algorithm was constrained to find no more than one change point per block of trials (AMOC method, “At Most One Change”) and using the Schwarz information criterion as the penalty term (default parameters for the cpt.mean function). The model was fit on the choice data over the 60 trials that comprised the blocks immediately preceding and following each action-outcome probability reversal, with the reversal centered at 0. Therefore, negative changepoint values indicated that the rat shifted behaviourally before the action-outcome contingencies reversed, while positive values indicated that they shifted their response after the reversal.

Choice latency was defined as the time between the first contact with the reward spout from the previous trial (or from the session start for the first trial), and the time of entry into the nosepoke. This time included the 0.5-second reward delivery on rewarded trials. However, rats typically remained near the spout to lick after the pump turned off, and stayed there for some time even on unrewarded trials. Collection latencies were defined as the time between the cue above the reward port and the first lick at the spout. Rats had to respond to the reward spout to begin a new trial regardless of reward delivery.

For blocked analysis, only the initial six blocks were included. The first cohort completed on average versus 6 blocks (t(16) = −17.141, p = 4.07e-41, independent t-test). The second cohort completed significantly more blocks. We therefore only analyzed the first six blocks across rats, with the goal of limiting bias on the performance measures by the second cohort. Importantly, we found no differences in performance between blocks blocks 4, 5 and 6, regardless of how many blocks were completed. However, there were sometimes decreases in trial completion rate and accuracy toward the end of the session and as the rats became satiated or less engaged. These periods of task performance were not included in any of the reported analyses.

### Reinforcement Learning

Four Q-learning models were fit to the choice behavior of the rats to estimate learning rates and inverse temperature. Two models were fit to Left/Right choice behavior while another two were fit to Stay/Shift choice behavior. For each type of choice behavior, we fit one model with a single learning rate and one with a separate learning rate for each option. Each model updated the value, *Q*, of a chosen option *i* based on reward feedback *r* as:

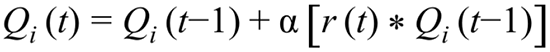

At each trial *t*, the updated value of an option *i* was given by its old value, *Q_i_(t−1)* plus a change based on the reward prediction error *(r(t)−Q_i_(t−1))*, multiplied by the learning rate parameter for that option, *α_i_*. In single-alpha models, *α_i_* referred to the same value for both options. Choice probability for each option *i* was calculated as *d_i_(t)* using:

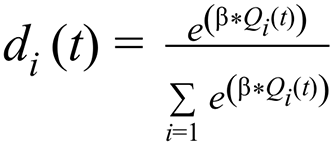

We then calculated the log-likelihood as:

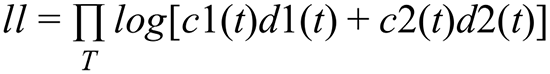

where *c_i_(t)* = 1 if the subject chooses option *i* in trial *t* and *c_i_(t)* = 0 otherwise. T is the total number of trials in the session or block. Parameters were fit by maximizing the log-likelihood using standard optimization techniques via the scipy package for python. Initial values for learning rate parameters were drawn from a normal distribution with a mean of 0.5 and a standard deviation of 0.5. Learning rate values were bounded within (0,1]. Initial values for the inverse temperature parameter were drawn from a normal distribution with a mean of 1 and a standard deviation of 5 and were bounded between 0 and 20. Initial values of actions were always set to 0.5 to minimize the impact of starting value on the other parameters by animal. When fitting blocks, the final action values of each block were carried over to initiate the next block. Model fits were repeated 100 times to avoid local minima and the model with the minimum negative log-likelihood was selected as the best fit. The Akaike Information Criterion (AIC) was then calculated for each best-fit model and was used to compare the goodness of fit between models. Because the number of free parameters differed by model, we also calculated the Bayesian Information Criterion (BIC), which more strongly penalizes a larger number of free parameters. While the best fit model according to both criteria had individual learning rates for each location, as previously found (Noworyta-Sokolowska et al., 2019), the learning rates were not statistically different from each other. Therefore, a simpler model with a single alpha parameter that updated both options was sufficient to model the behavioral data here.

## Results

### Effects of Reward Schedule on Performance

We trained 17 rats on a spatial, blocked two-armed bandit task. The rats were first trained deterministically (100%/0%), and then tested on 90%/10%, 80%/20% and 70%/30% probabilistic reward schedules. Each schedule was repeated one session per day for three days. The rats had to nosepoke one of two options on one side of the box. They immediately received tonal feedback as to whether that choice was rewarded. Regardless of the outcome, the rat had to lick at a spout on the opposite side of the box. The spout delivered a reward on reinforced trials or immediately started the next trial on non-rewarded trials. Reward probabilities reversed every 30 trials (Figure 1A). We trained the rats in two groups. The second cohort of rats (n=11) was trained to a minimum accuracy criterion as set by the first cohort (n=6). For each session, we calculated the proportion of Win trials in which the animals stayed with the same port (Win-Stay) and the proportion of Loss trials in which the animals shifted to the other port (Lose-Shift) (Figure 1B), but this measure was not used as a criterion for training.

**Figure 1.**
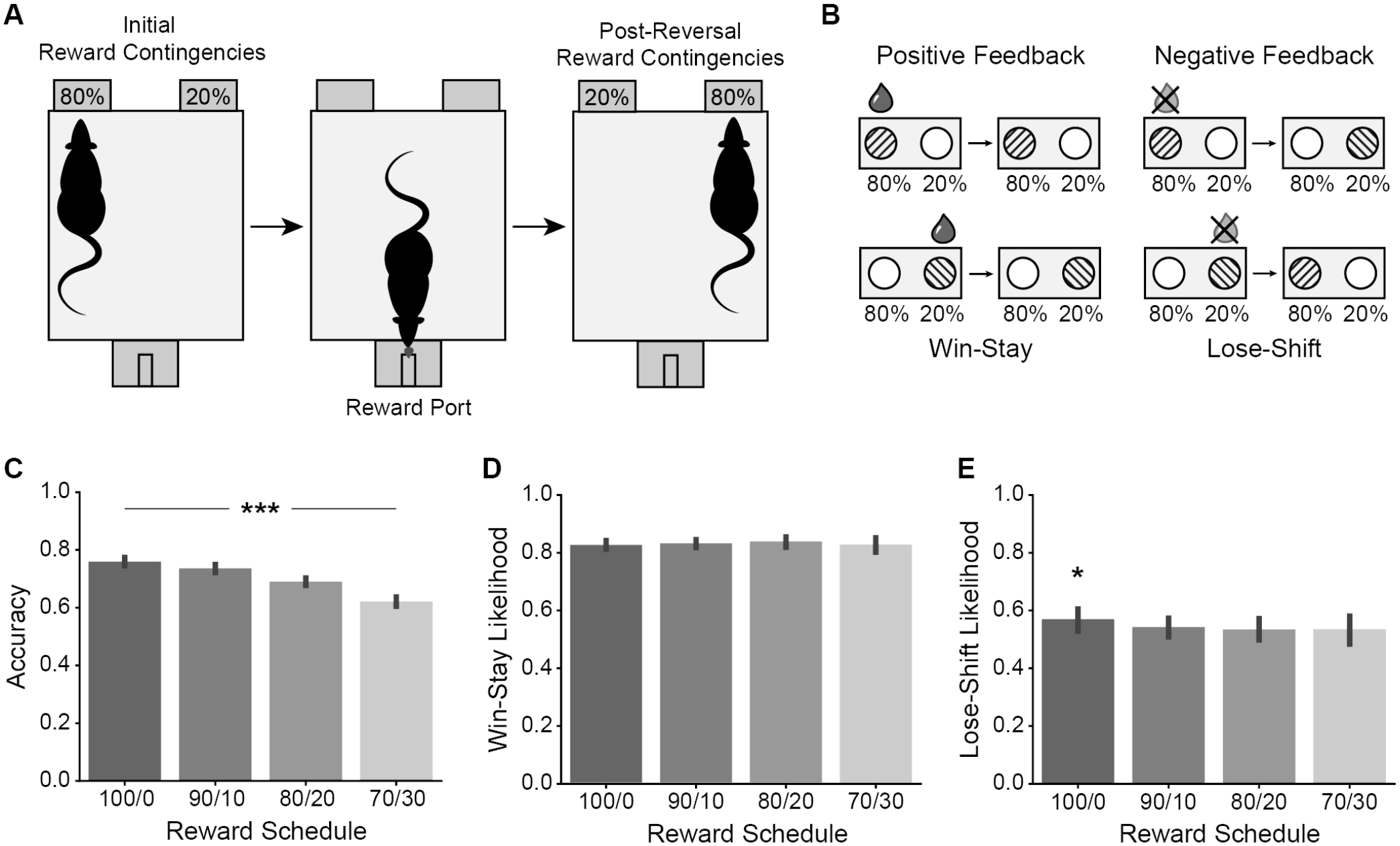
Design and Performance of the Two-Armed Bandit Task. (A) Task design: Rats were presented with two nosepokes on one side of the chamber. Each nosepoke was associated with a different probability of reward delivery. The reward spout was located on the opposite side of the chamber. The contingency between nosepoke and reward probability reversed every 30 trials throughout the session. (B) Win-Stay/Lose-Shift strategies: Each trial can be described as a reaction to the feedback from the previous trial. Win-Stay decisions occur when the same option is repeated following positive feedback. Lose-Shift trials occur when the alternative option is selected following negative feedback. (C) Reward schedule impacted accuracy across three days of testing per schedule (F(3,46) = 92.162, p = 0, ANOVA). (D) Reward schedule had no influence on mean Win-Stay likelihood (F(3,46) = 0.971, p = 0.415, ANOVA). (E) Reward schedule was shown to influence mean Lose-Shift likelihood (F(3,46) = 2.492, p = 0.0719, ANOVA), and this effect was driven by a higher LS likelihood under the deterministic versus probabilistic schedules (p = 0.0104, post-hoc Tukey test). Error bars denote 95% confidence intervals.

Accuracy, defined as the percentage of trials targeted toward the higher-value option, varied significantly with reward schedule (F(3,46) = 92.162, p = 0, ANOVA, Figure 1C). In the deterministic version of the task, when feedback was most informative, subjects performed well (mean accuracy = 77.83%). Subjects performed more poorly when the reward feedback was less revealing of the higher-value target. For example, at the 70/30 reward schedule, rats were only able to obtain a mean accuracy of 63.09% over a session.

We also note that the number of blocks completed varied by rat (3-24 blocks, mean = 11), and the second cohort completed significantly more blocks than the first, 15 blocks on average versus 6 blocks (t(16) = −17.141, p = 4.07e-41, independent t-test). Accuracy was lower in the second cohort in general (t(16) = 2.7546, p = 0.00642, independent t-test), but there was no effect of the number of blocks completed on accuracy (t(16) = −1.379, p = 0.169, linear regression). There was no effect of cohort for any other measure. There was also no effect of day over the 3-day test for each reward schedule (F(2,30) = 2.548, p = 0.0951, ANOVA), though there was a slight interaction between schedule and day (F(6,94) = 2.646, p = 0.0205, ANOVA).

To determine whether the decrease in accuracy was due to a change in sensitivity to positive and negative feedback, we analyzed their Win-Stay/Lose-Shift (WSLS) decision strategy for each schedule. Their overall sensitivity to positive and negative feedback was approximated by determining the proportion of WS and LS trials in a session. There was no difference by reward schedule in Win-Stay likelihood (F(3,46) = 0.971, p = 0.415, ANOVA, Figure 1D) and no effect of day within schedule (F(1,46) = 0.054, p = 0.818, ANOVA). There was of reward schedule on Lose-Shift likelihood (F(3,46) = 2.492, p = 0.0719, ANOVA, Figure 1E). These results indicate that the rats did not adjust their strategy to compensate for reward uncertainty beyond this level.

### Effects of Block on Performance

Since the task required many spatial reversals within the session, it is possible that the rats’ strategy, and therefore performance, changed over time. To investigate within-session trends, we independently analyzed each block. A repeated-measures ANOVA confirmed the effect of block on accuracy (F-stat(5,74) = 3.34, p = 0.00895, ANOVA, Figure 2A). Subjects demonstrated a drop in accuracy in the second block (following the first reversal) in all reward schedules (p = 0.0000304, post-hoc Tukey test). There was no interaction between block and schedule (F(5,74) = 0.754, p = 0.586, ANOVA).

**Figure 2.**
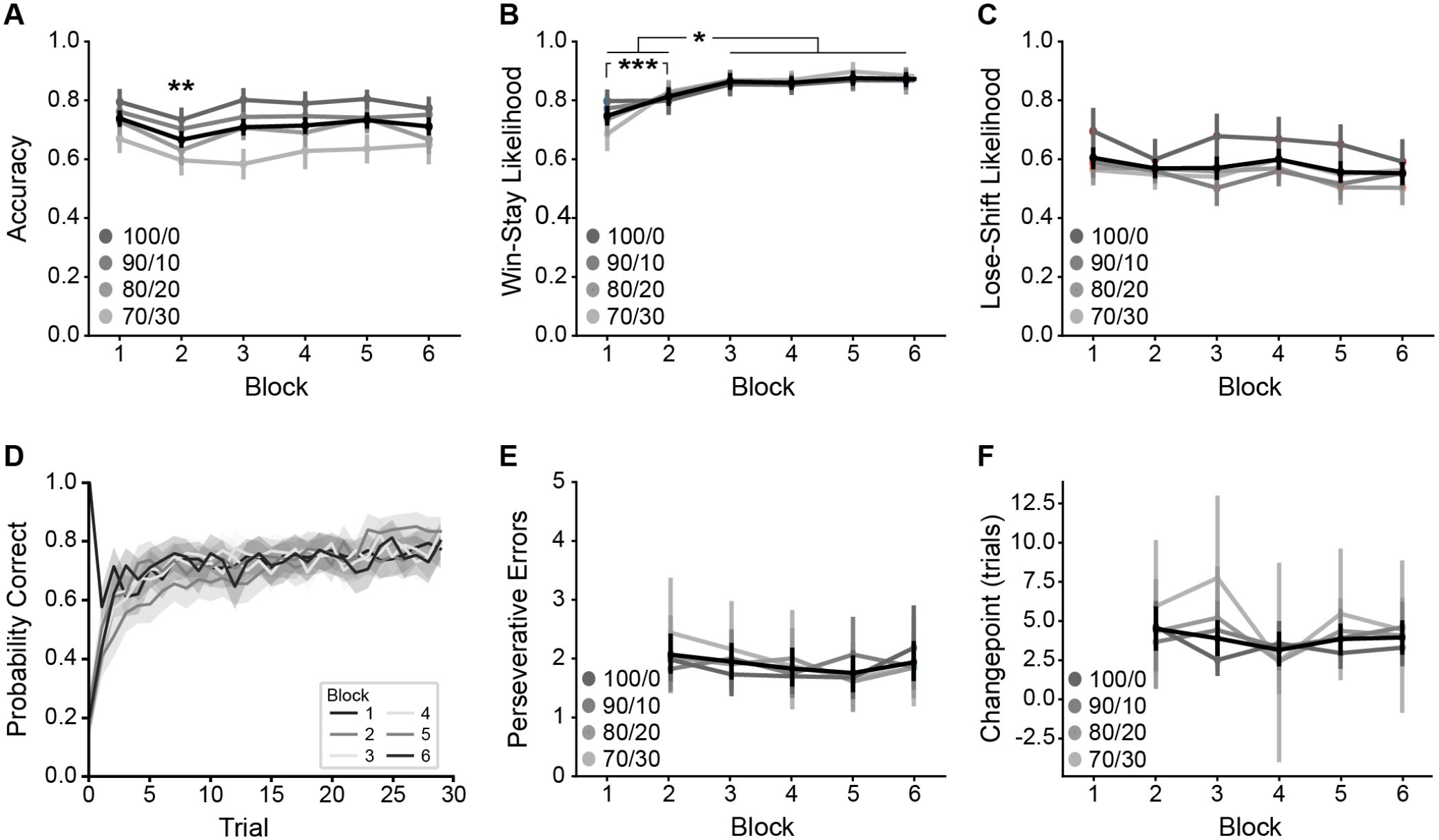
Behavior Over Blocks in the Two-Armed Bandit Task. (A) Block number had an effect on mean accuracy early in the session (F-stat(5,74) = 3.34, p = 0.00895, ANOVA). Specifically, there was a difference between the first and second blocks in all reward schedules (p = 0.0000304, post-hoc Tukey test). (B) Average WS likelihood increased across blocks early in the session. F(5,74) = 23.562, p < 0, ANOVA). The first and second block were significantly lower than the subsequent four blocks (p < 0.0249, post-hoc Tukey test), and were different from each other (p = 0.000152, post-hoc Tukey test). (C) Although an ANOVA reported a block-dependent change in mean LS likelihood (F(5,74) = 3.056, p = 0.0146, ANOVA), post-hoc testing did not indicate a difference between any two specific blocks. (D) The average probability of selecting the “correct” option increased across trials. In the second block, this probability increased comparatively slowly. (E) There was no difference in the mean number of perseverative errors following each reversal (F(4,64) = 0.593, p = 0.669, ANOVA). F) Reward schedule had no impact on mean changepoint (F(1,33) = 2.383, p = 0.132, ANOVA) and there was no within-session effect of block (F(4,33) = 2.037, p = 0.112, ANOVA). Significance: * p < 0.05, ** p < 0.01, *** p < 0.001. Error bars denote 95% confidence intervals.

Within the shortened view of the first six blocks, there was still no difference in WS likelihood by reward schedule (F(1,74) = 1.646, p = 0.204, ANOVA), but there was a difference across blocks (F(5,74) = 23.562, p = 0, ANOVA, Figure 2B). Post-hoc testing revealed that the first and second block were significantly lower than the other four blocks (p < 0.0249, post-hoc Tukey test), and also that they were different from each other (p = 0.000152, post-hoc Tukey test). Repeated-measures ANOVA also reported a block-dependent change in LS likelihood(F(5,74) = 3.056, p = 0.0146, ANOVA, Figure 2C), but post-hoc testing did not indicate a difference between any two specific blocks. There was also no interaction between block and schedule (F(5,74) = 0.929, p = 0.467, ANOVA). Together, these results indicate that the strategy for positive feedback changes with experience and eventually collapses to some value across all reward schedules, while the strategy for negative feedback depends on whether the task is probabilistic but does not change with experience.

We expected that increased uncertainty might also cause perseveration around reversals, since the animals would have more difficulty distinguishing between probabilistic negative feedback and a true reversal, and because their Lose-Shift rates were lower in probabilistic schedules. However, there was no statistical difference in the number of perseverative errors by reward schedule (F(3,48) = 1.444, p = 0.242, ANOVA). Interestingly, their WS likelihood was lowest in the first block, while their accuracy was lowest in the second. Additionally, the mean likelihood of selecting the correct side by trial appeared to reach asymptote in the second block more slowly than for the other blocks (Figure 2D). To investigate the decrease in accuracy in the second block, we attempted to quantify how quickly rats adapted to each reversal. However, rats made the same number of perseverative errors following each of the first five reversals (F(4,64) = 0.593, p = 0.669, ANOVA, Figure 2E). The mean number of perseverative errors was 1.78 trials.

It is possible that while they always chose the target option within a few trials after a reversal, it took longer for the rats to reliably choose the target option within the block. To estimate this behavioral reversal for each block, we applied a changepoint algorithm to the rats’ choice data (see Methods). Reward schedule had no impact on behavioral changepoint (F(1,33) = 2.383, p = 0.132, ANOVA). There was also no within-session effect of block (F(4,33) = 2.037, p = 0.112, ANOVA, Figure 2F), however the algorithm was only able to detect a statistically valid changepoint after 57% of the first 5 reversals in each session. Out of those, the rats demonstrated a behavioral reversal 3.32 trials after the environmental reversal on average. This finding suggests that the rats shifted their behavior abruptly soon after the reversal in action-outcome contingencies for half of the blocks, but for the other half of blocks gradually learned the new contingencies or did not demonstrate a behavioral reversal at all.

### Reinforcement Learning Analysis

WSLS strategies describe the overall trends in individual choices but may not describe behavior over longer time periods adequately, as indicated by the change in WS likelihood by block. We fit several reinforcement learning models to the data to determine if such a model that incorporates outcome history would better account for their behavior. We fit the model using either left-right or stay-shift dichotomies as the options to learn, and tried models with either a single learning rate or with separate learning rates for each option. The model that had the lowest Akaike Information Criterion (AIC) used individual learning rates to update the values of left and right actions. However, since there was no statistical difference between the alpha values for this model (t(201) = 0.2659, p = 0.7906, paired t-test), there was no real difference in the learning rate for each option and a better fit is likely due to the freedom given by the extra parameter. Therefore, we elected to use the single-alpha model instead.

Uncertainty affected learning rate (alpha) in the single-alpha model (F(3, 46) = 3.591, p = 0.0205, ANOVA, Figure 3A). Post-hoc testing showed that the 70/30 schedule specifically produced slower learning, driving the effect (p < 0.09, post-hoc Tukey test). There was no corresponding effect on inverse temperature (beta) (F(3,46) = 0.77, p = 0.517, ANOVA). Log transforms of the values also found no effect (F(3,46) = 1.022, p = 0.392. However, there were differences in variance in the fits of beta between all pairs of reward schedules with the exception of 100/0 - 80/20 (F(1,49), p < 0.000148, variance test), which violated the assumptions of the ANOVA. Nonparametric testing revealed dramatic differences in the medians of the fits (χ^2^(3) = 71.976, p = 1.611e-15, Kruskal-Wallis rank sum test, Figure 3B), such that beta decreased with uncertainty.

**Figure 3.**
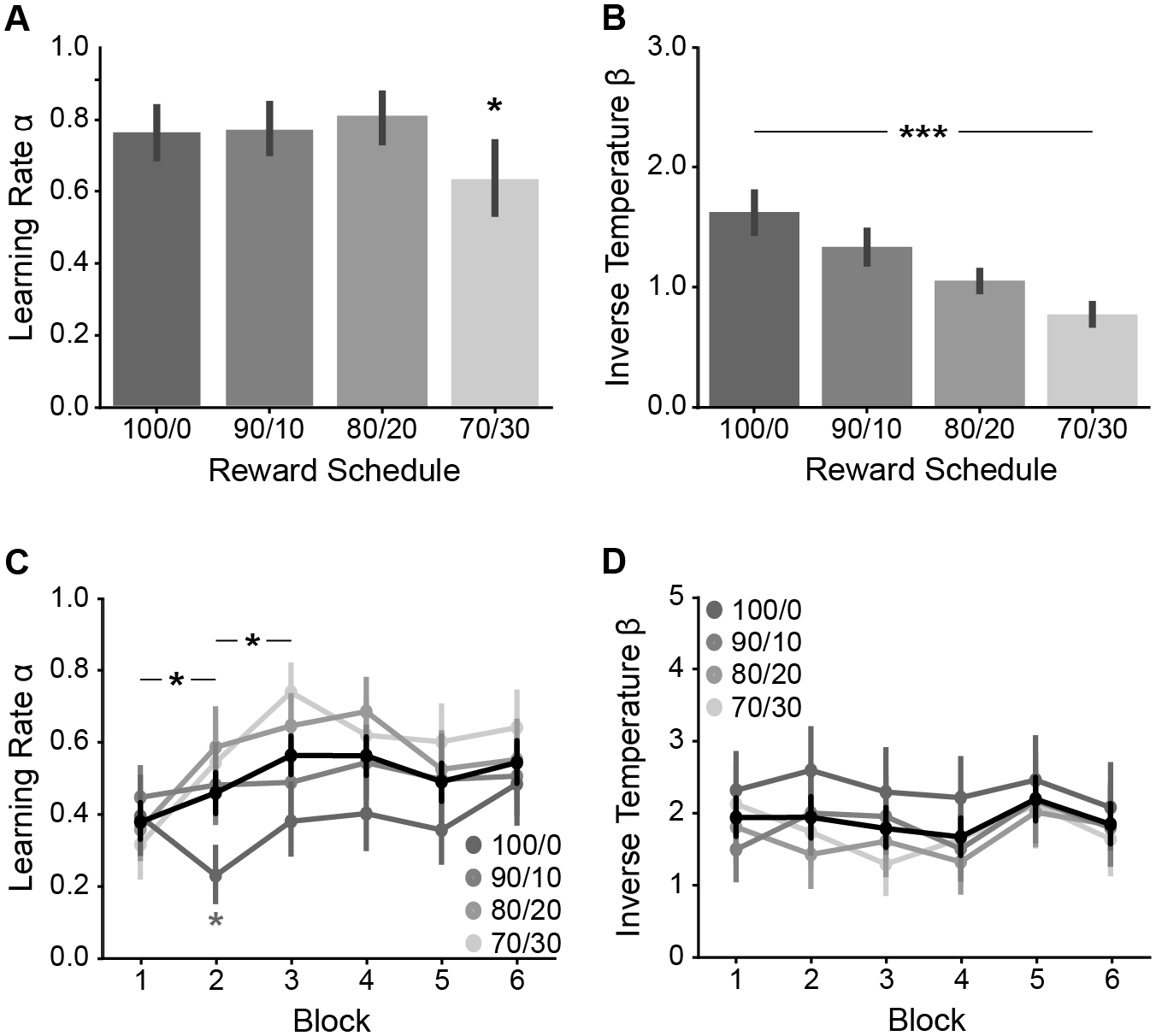
Modelling of Behavior with Reinforcement Learning. (A) Learning rate was affected by reward schedule (F(3, 46) = 3.591, p = 0.0205, ANOVA), and the effect was specifically caused by slower learning under the most difficult 70/30 schedule (p < 0.09, post-hoc Tukey test). (B) The median inverse temperature across rats decreased with increasing uncertainty (χ^2^(3) = 71.976, p = 1.611e-15, Kruskal-Wallis rank sum test). (C) An additional RL model was fit to each of the first six blocks individually and found evidence that learning rate varied by block (F(5,65) = 9.321, p = 9.92e-07, ANOVA). (D) There was no effect of block on mean inverse temperature (F(5,65) = 1.543, p = 0.174, ANOVA). Significance: * p < 0.05, ** p < 0.01, *** p < 0.001. Error bars denote 95% confidence intervals.

One of the reasons we used a blocked design was that we could apply reinforcement learning to each block individually. When we fit the model to each of the first six 30-trial blocks, reward schedule impacted both learning rate (F(3,65) = 4.152, p = 0.00938, ANOVA) but not inverse temperature (F(3,65) = 2.064, p = 0.114, ANOVA). The effect of learning rate was dominated by model parameters for the deterministic 100/0 schedule (p < 0.0024, post-hoc Tukey test). Learning rate also varied by block regardless of schedule, (F(5,65) = 9.321, p = 9.92e-07, ANOVA, Figure 3C). Within the probabilistic schedules, learning rate increased over the first three blocks. There was no effect of block on inverse temperature (beta parameter) (F(5,65) = 1.543, p = 0.174, ANOVA, Figure 3D) and no interaction with reward schedule (F(12,65) = 1.442, p = 0.170, ANOVA). When fit to each block individually, we found homoscedasticity (equal variance) across reward schedules. Nevertheless, to confirm these results using a non-parametric method, we found no difference in median beta by block (χ2(5) = 9.38, p = 0.0948, Kruskal-Wallis rank sum test).

In the deterministic session, learning rate drops in the second block and recovers in the third block, but this was not found to be significant in a post-hoc test. In contrast, there was no such effect of block on inverse temperature (beta parameter) (F(5,65) = 1.543, p = 0.174, ANOVA, Figure 3D) and no interaction with reward schedule (F(12,65) = 1.442, p = 0.170, ANOVA). When fit to each block individually, we found homoscedasticity (equal variance) across reward schedules. Nevertheless, to confirm these results using a non-parametric method, we found no difference in median beta by block (χ^2^(5) = 9.38, p = 0.0948, Kruskal-Wallis rank sum test).

To quantify the relationship between WSLS strategy and RL parameters, we performed a linear regression on the WSLS values to predict learning rate and inverse temperature. Both alpha (t(3,198) = 2.461, p = 0.0147, linear regression) and beta (t(3,198) = −2.847, p = 0.00488, linear regression) were best predicted by the interaction between Win-Stay and Lose-Shift likelihoods. For each rat, LS (sensitivity to negative feedback) was a better predictor of learning rate (WS: t(3,198) = −1.890, p =0.0602; LS: t(3,198) = −2.246, p = 0.0258) and WS (sensitivity to positive feedback) was a better predictor of choice stochasticity (WS: t(3,198) = 2.777, p = 0.00602; LS: t(3,198) = 2.714, p = 0.00724).

### Inactivation of Medial Orbitofrontal Cortex

To determine the role of mOFC in the blocked TAB design, a subset of rats (n=9) were implanted with cannula bilaterally in mOFC and received infusions of muscimol one hour prior to testing (Figure 4A). Because both accuracy and LS likelihood were highest in the 100/0 session, all rats were tested in the deterministic reward schedule only to increase the likelihood of detecting an effect and to limit risk associated with intra-cortical infusion. Inactivation of mOFC resulted in a decrease in accuracy (F(1,8) = 7.979, p = 0.0223, ANOVA, Figure 4B). While inactivation did not affect Win-Stay likelihood (F(1,8) = 0.085, p = 0.778, ANOVA, Figure 4C), it caused a decrease in Lose-Shift likelihood (F(1,8) = 12.53, p = 0.00763, ANOVA, FIgure 4D). This decrease in LS likelihood corresponded to an increase in changepoint (F(1,14) = 9.695, p = 0.00762, ANOVA, Figure 4E) and perseverative errors (F(1,32) = 5.513, p = 0.0252, ANOVA, Figure 4F).

**Figure 4.**
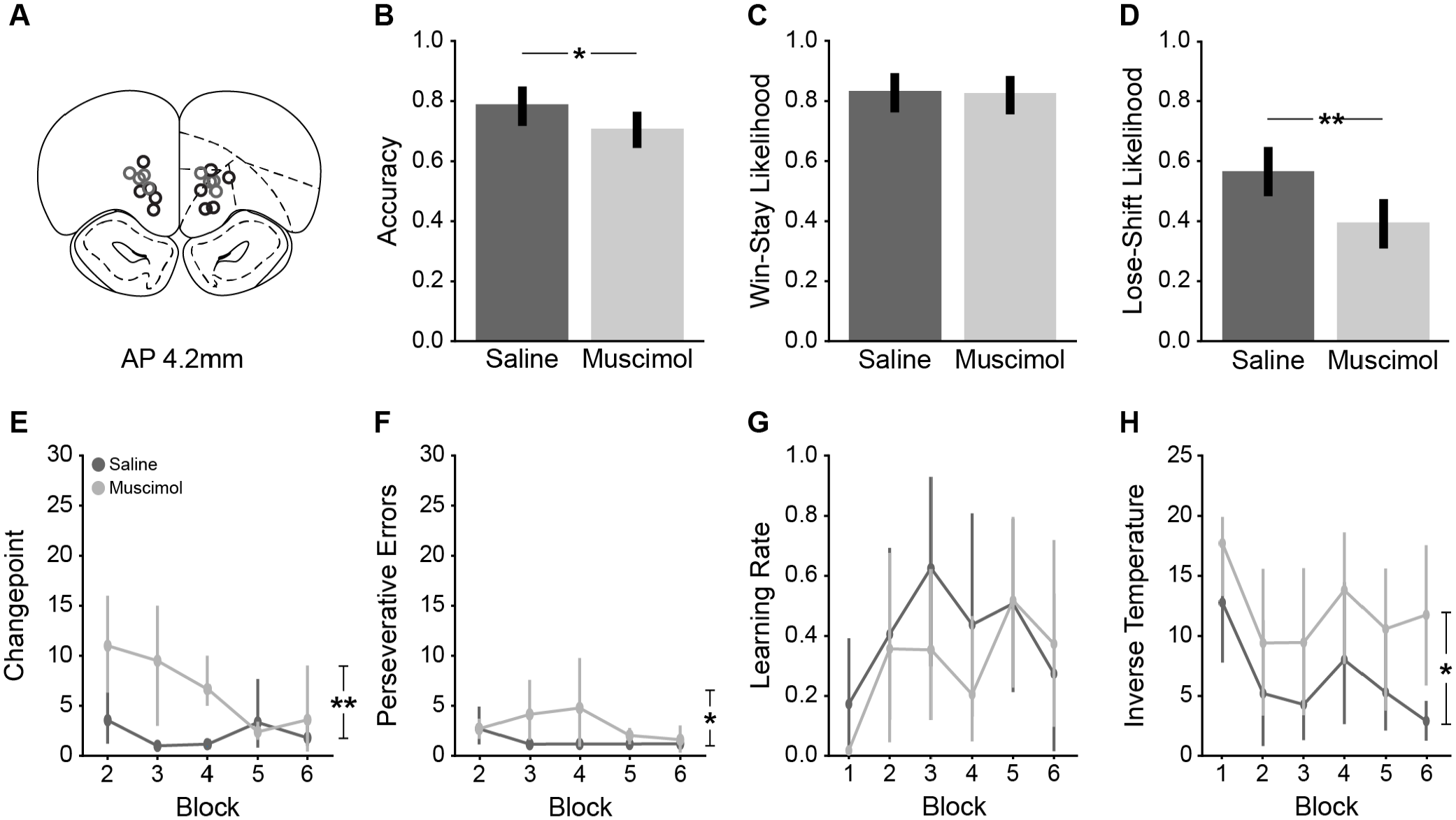
Inactivation of mOFC Decreased Sensitivity to Negative Feedback. (A) Bilateral guide cannulae were implanted in mOFC and all animals were tested under muscimol (reversible inactvation). Infusion locations highlighted in blue were also tested with intra-cortical yohimbine. (B) Inactivation of mOFC resulted in a decrease in accuracy (F(1,8) = 7.979, p = 0.0223, ANOVA). (C) Inactivation did not affect Win-Stay likelihood (F(1,8) = 0.085, p = 0.778, ANOVA). (D) Lose-Shift likelihood decreased following mOFC inactivation (F(1,8) = 12353, p= 0.00763, ANOVA). (E) Inactivation increased changepoint (F(1,14) = 9.695, p = 0.00762, ANOVA). (F) There was a slight increase in perseveration following mOFC inactivation (F(1,32) = 5.513, p = 0.0252, ANOVA). (G) Inactivation of mOFC did not impact learning rate (F(1,16) = 0.553, p = 0.468). There was also no effect of block (F(5,32) = 0.868, p = 0.513). (H) Inactivation of mOFC did not affect inverse temperature (χ^2^(1) = 0.43905, p-value = 0.5076). However, there was a difference in inverse temperature when the RL model was fit by block (F(1,5) = 13.461, p = 0.0145, ANOVA), with muscimol increasing the beta parameter. This effect did not interact with block (F(1,5) = 0.134, p = 0.587, ANOVA). Significance: * p < 0.05, ** p < 0.01, *** p < 0.001. Error bars denote 95% confidence intervals.

Inactivation of mOFC did not affect either the learning rate (F(1,8) = 2.709, p = 0.138) or inverse temperature (χ^2^(1) = 0.43905, p = 0.5076) in a session-wide analysis of reinforcement learning. There was no interaction with learning rate when analyzed by block (F(5,32) = 0.868, p = 0.513). However, there was a difference in inverse temperature when the RL model was fit by block (F(1,5) = 13.461, p = 0.0145, ANOVA), with muscimol increasing the beta parameter. There was no interaction between treatment and block (F(1,5) = 0.336, p = 0.587, ANOVA).

### Effects of Systemic Yohimbine

To investigate the role of norepinephrine in two-armed bandit performance, we first challenged a subset (n=11) of the rats with a 2 mg/kg systemic injection of yohimbine. They were tested on both deterministic and 80/20 probabilistic reward schedules since IP injections posed less of a health risk than intra-cortical infusion. If norepinephrine moderates the balance between exploration and exploitation, we would expect to see a change in their strategy. Indeed, there was a decrease in WS likelihood in both reward schedules under yohimbine (F(1,10) = 21.43, p = 0.000936, ANOVA, Figure 5C). There was no change in LS likelihood (F(1,10) = 2.078, p = 0.18, ANOVA, Figure 5D), but as before, there was a significant difference in LS strategy between the 100/0 and 80/20 in the saline session (F(1,10) = 7.722, p = 0.0195, ANOVA). Therefore, yohimbine selectively decreased sensitivity to positive feedback.

**Figure 5.**
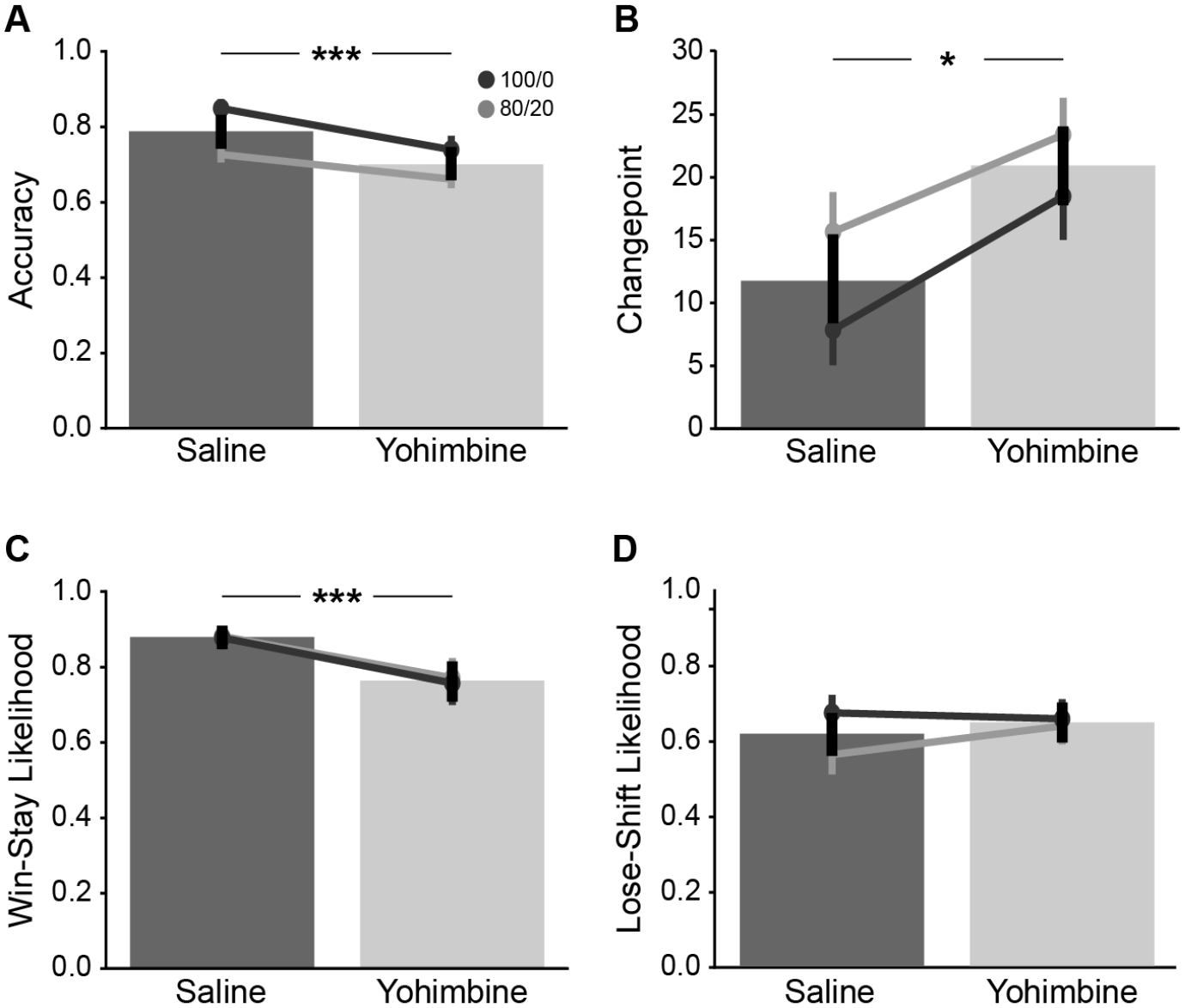
Systemic Yohimbine Reduced Sensitivity to Positive Feedback. Rats were challenged with a 2 mg/kg systemic injection of yohimbine or saline control and tested under deterministic (dark line) and 80/20 probabilistic (light line) reward schedules. (A) Yohimbine decreased accuracy compared to saline controls (F(1,10) = 29.03, p = 0.000306, ANOVA). (B) Yohimbine increased changepoint compared to saline controls (F(1,124) = 4.664, p = 0.03273, ANOVA). (C) Average WS likelihood decreased in both reward schedules under yohimbine as compared to control sessions (F(1,10) = 21.43, p = 0.000936, ANOVA). (D) There was no change in mean LS likelihood under yohimbine (F(1,10) = 2.078, p = 0.18, ANOVA). Significance: * p < 0.05, ** p < 0.01, *** p < 0.001. Error bars denote 95% confidence intervals.

This effect led to a significant decrease in accuracy under yohimbine compared to saline control (F(1,10) = 29.03, p = 0.000306, ANOVA, Figure 5A). As with the uncertainty testing, animals were more accurate in the deterministic reward schedule (F(1,10) = 54.9, p = 2.29e-05, ANOVA). However, there was no interaction between treatment and schedule (F(1,10) = 3.416, p = 0.0943, ANOVA), meaning that yohimbine caused a similar decrease in accuracy in both tests. This decrease in accuracy was not associated with perseveration, as treatment with yohimbine did not impact the number of perseverative errors the animals made following each reversal (F(1,10) = 0.925, p = 0.359, ANOVA). However, yohimbine did increase the behavioral changepoint (F(1,10) = 9.497, p = 0.0116, ANOVA, Figure 5B).

We then focused on the first 6 blocks to investigate yohimbine’s effect on the first few reversals. While treatment decreased the likelihood of choosing the correct option, there was no interaction between treatment and block on accuracy (F(5,50) = 1.468, p = 0.217, ANOVA), on Win-Stay likelihood (F(5,50) = 1.808, p = 0.128, ANOVA), or Lose-Shift likelihood (F(5,50) = 0.766, p = 0.579, ANOVA), at least in the context of the order of the blocks. There was however a difference in accuracy between even and odd blocks for the deterministic session, (t(10) = 4.275, p = 6.394e-05, dependent t-test, Figure 6A) with lower accuracy on the odd blocks. This suggests that the rats become biased toward the initial high-value option. This effect was present in Win-Stay likelihood (t(10) = 3.141, p = 0.0025, dependent t-test, Figure 6B), but there was no such pattern in LS likelihood (t(10) = 0.260, p = 0.795, dependent t-test, Figure 6C). However, there was no significant effect on accuracy for the 80/20 probabilistic schedule (t(10) = 0.256, p = 0.799, dependent t-test), suggesting that the effects of yohimbine on bias depend on reward uncertainty.

**Figure 6.**
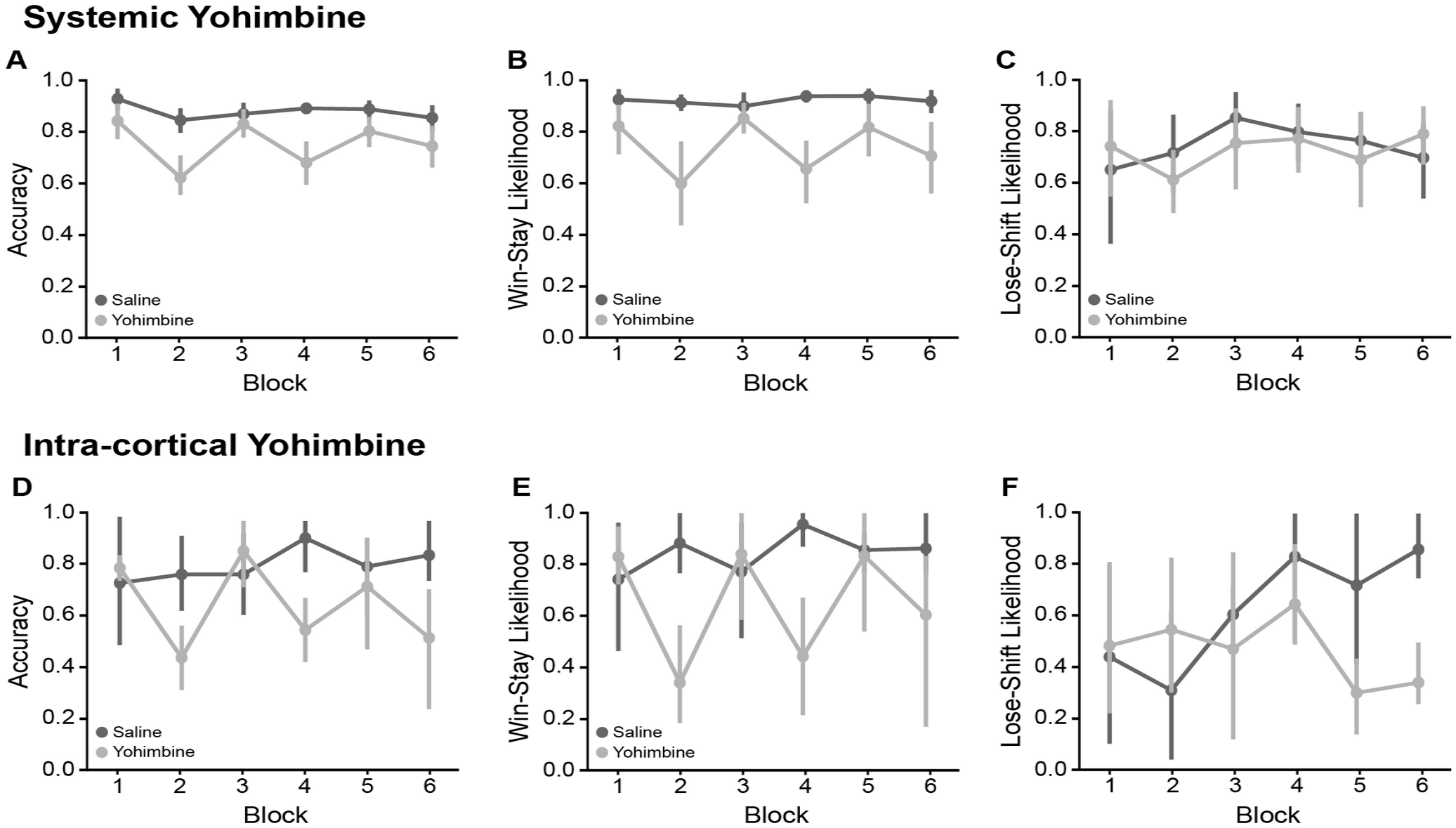
Systemic yohimbine altered choice behavior under a deterministic reward schedule. (A) Rats were more accurate in odd blocks, compared to even blocks in sessions with systemic yohimbine (t(10) = 4.275, p = 6.394-05, dependent t-test). (B) Rats were more likely to Win-Stay on odd blocks compared to even blocks (t(10) = 3.141, p = 0.0025, dependent t-test). (C) There was no difference in LS likelihood between even and odd blocks (t(10) = 0.260, p = 0.795, dependent t-test). (D) Intra-cortical yohimbine did not change mean accuracy compared to saline control on even blocks (t(3) = 1.958, p = 0.057, independent t-test). (E) Intra-cortical yohimbine did not affect WS likelihood on even blocks (t(3)= 1.789, p = 0.081, independent t-test). (F) Intra-cortical yohimbine did not affect LS likelihood (t(3) = −0.765, p = 0.45, independent t-test). Significance: * p < 0.05, ** p < 0.01, *** p < 0.001. Error bars denote 95% confidence intervals.

There was also a double dissociation of the effect by reward schedule and block under yohimbine (F(5,120) = 3.405, p = 0.00652, ANOVA). Under the deterministic schedule yohimbine increased the number of trials the rats needed to behaviorally reverse for each of the first six reversals. Under the probabilistic schedule however, they reversed their behavior before the first two environmental reversals in the session, and then took longer to adapt after the next three reversals than in the saline session.

We predicted that norepinephrine controls the determinism of choice via control over the inverse temperature in reinforcement learning. When we fit the single-alpha reinforcement learning model discussed above to their behavior to the whole session, there was no change in learning rate under yohimbine (F(1,10) = 2.162, p = 0.172, ANOVA, Figure 7A). In contrast, we found that yohimbine decreased inverse temperature as predicted (F(1,10) = 6.31, p = 0.0308, ANOVA, Figure 7B). There was also no interaction with schedule for either parameter (Alpha: F(1,10) = 0.295, p = 0.599; Beta: F(1,30) = 0.83, p = 0.384, ANOVA). These fits had equal variance and did not require non-parametric analysis.

**Figure 7.**
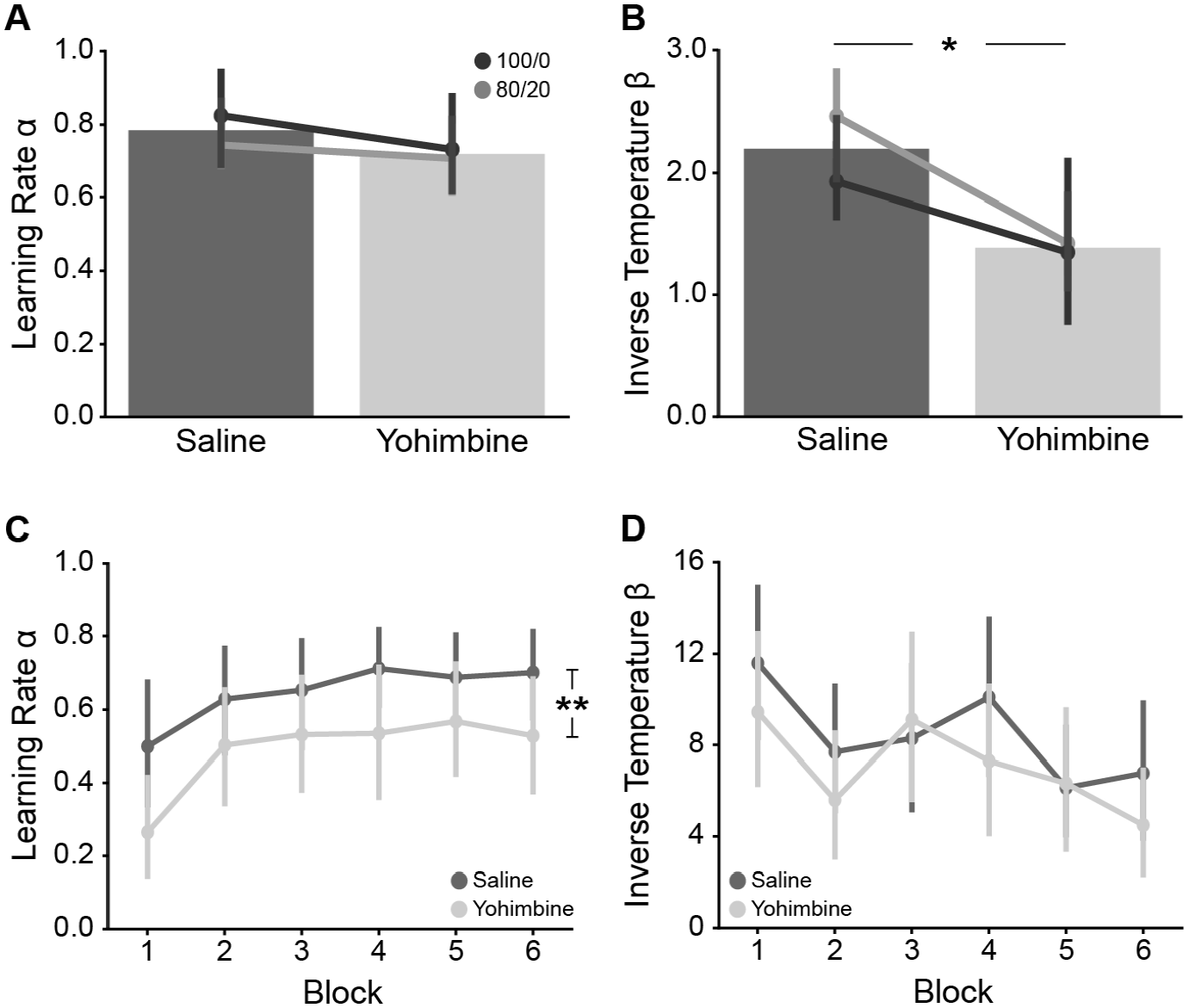
Yohimbine Altered Learning Rate and Inverse Temperature. (A) Systemic yohimbine did not affect learning rate, measured over the entire session (F(1,10) = 2.162, p = 0.172, ANOVA). (B) Yohimbine decreased inverse temperature, measured over the entire session (F(1,10) = 6.31, p = 0.0308, ANOVA). (C) When the model was fit by block, yohimbine decreased learning rate (F(1,10) = 10.06, p = 0.00995, ANOVA). There was an effect of block on learning rate (Alpha: F(5,50) = 2.583, p = 0.0373, ANOVA), but no interaction with treatment (F(5,50) = 1.275, p = 0.289, ANOVA). (D) When the model was fit by block, yohimbine did not affect inverse temperature (F(1,10) = 3.067, p = 0.11, ANOVA). There was an effect of block (F(5,50) = 2.942, p = 0.021, ANOVA), but no interaction between treatment and block (F(5,50) = 1.421, p = 0.233, ANOVA). Significance: * p < 0.05, ** p < 0.01, *** p < 0.001. Error bars denote 95% confidence intervals.

When fit to each block individually, we found that yohimbine treatment decreased learning rate (F(1,10) = 10.06, p = 0.00995, ANOVA, Figure 7C) but not inverse temperature (F(1,10) = 3.067, p = 0.11, ANOVA, Figure 7D). While there was an effect of block for both parameters(Alpha: F(5,50) = 2.583, p = 0.0373; Beta: F(5,50) = 2.942, p = 0.021, ANOVA), there was no interaction between treatment and block (Alpha: F(5,50) = 0.226, p = 0.949; Beta: F(5,50) = 0.469, p = 0.797, ANOVA). Because there was no interaction with schedule (Alpha: F(5,50) = 1.275, p = 0.289; Beta: F(5,50) = 1.421, p = 0.233, ANOVA), we combined the results of the fits in Figure 7C-D.

### Intra-cortical Infusions of Yohimbine

A subset of rats (n=4) were also bilaterally infused with yohimbine in medial OFC under the deterministic schedule (see Figure 4A for this schedule) to directly compare against the muscimol infusion above. Blockade of α2 adrenergic receptors in this brain region caused a similar reduction in accuracy (F(1,3) = 12.74, p = 0.0376, ANOVA), and Win-Stay Likelihood (F(1,3) = 16.68, p = 0.0265, ANOVA) across the session, and did not affect Lose-Shift likelihood (F(1,3) = 0.079, p = 0.796, ANOVA). Again, as with the systemic injections, yohimbine did not affect the number of perseverative errors (F(1,3) = 0.247, p = 0.653, ANOVA), nor did it increase changepoint (F(1,3) = 7.321, p = 0.0734, ANOVA) in these rats. When analyzed by even versus odd blocks, we found no effect on accuracy (t(3) = 1.958, p = 0.057, independent t-test, Figure 6D) or WS likelihood (t(3)= 1.789, p = 0.081, independent t-test, Figure 6E). There was also no such effect on LS likelihood (t(3) = −0.765, p = 0.45, independent t-test, Figure 6F). The effects of reinforcement learning did not reach the threshold for statistical significance, due to the small group size and high variability of model fits between rats.

### Effects of Yohimbine on Choice and Reward Collection Latencies

We hypothesized that rats might take longer to make decisions when feedback is less informative. However, there was no difference in median choice latency (F(3,30) = 1.13, p = 0.353, ANOVA) or collection latency (F(3,30) = 1.584, p = 0.214, ANOVA) by schedule. There was however a significant effect of block on both median choice latency (F(5,44) = 9.512, p = 3.34e-06, ANOVA) and collection latency (F(5,44) = 8.269, p = 1.42e-05, ANOVA). Specifically, rats had longer latencies in the initial block relative to all other blocks (Choice: t(10) = 3.4489, p = 0.0013; Collect: t(10) = 3.1982, p =0.0026, paired t-test). There was no interaction between block and schedule for either measure.

Systemic yohimbine decreased both choice latency (F(1,10) = 12.14, p = 0.00588, ANOVA) and collection latency (F(1,10) = 5.983, p = 0.0345, ANOVA) in the first six blocks. Block number affected choice latency in both sessions (F(5,50) = 5.375, p = 0.000496, ANOVA) with no interaction with drug treatment (F(5,50) = 2.326, p = 0.0881, ANOVA). The effect of block was even more pronounced for collection latency (F(5,50) = 7.999, p = 1.34e-05, ANOVA), as was the interaction with yohimbine (F(5,50) = 3.596, p = 0.00743, ANOVA).

In particular, rats were slower in the first block as compared to the second (Choice: p= 0.0251, Collect: p=0.00163, post-hoc Tukey test). There was also an overall increase in the mean number of blocks completed: 14.6 under systemic yohimbine vs 12.1 under the saline control, (t(43) = −4.4476, p = 0.0002, paired t-test), indicating that the increase in pace enabled them to complete more trials in the 1-hour session.

To examine this issue, we regressed median choice latency, treatment, and correctness onto accuracy. While drug treatment was a predictive factor as expected (β = - 2.756, p = 0.0075, linear regression), choice latency had no relationship with accuracy (β = - 0.348, p = 0.7290, linear regression) nor was the interaction between choice latency and systemic yohimbine (β = 0.590, p = 0.5569, linear regression).

In the four rats with guide cannula, intra-cortical yohimbine affected neither choice (F(1,3) = 0.287, p = 0.629) nor collection latencies (F(1,3) = 0.516, p = 0.524). However, it is unclear if this finding was due to the small sample size (n=4) or if these individual rats were unaffected, since systemic yohimbine also did not alter latencies in these animals (Choice: F(1,3) = 1.748, p = 0.278; Collect: F(1,3) = 0.199, p = 0.686).

## Discussion

Win-Stay Lose-Shift strategy has been shown to be consistent in rats across sessions in an 80/20 probabilistic bandit (Noworyta-Sokolowska et al., 2019), though further evidence shows that WS and LS likelihoods change in the learning and reversal phases of a single probabilistic reversal (Amodeo et al., 2017). In contrast, human choices may be best fit by a WSLS model that changes with experience (Worthy & Maddox, 2014). However, these studies confound WS likelihood with the reversal trigger. The current study used a blocked bandit and corroborates both findings to some extent. We found no change in LS likelihood by block, but likelihood of Lose-Shifting was higher in deterministic schedules as compared to probabilistic schedules. This is unsurprising as negative feedback is perfectly informative of a reversal in the deterministic schedule. In contrast, WS likelihood increased over the first three blocks but was stable across schedules. This finding is in contrast to a recent study that found increased Win-Shift behavior at the beginning of each block and in the higher uncertainty schedule within a 3-armed bandit (Cinotti et al., 2019). However, this study averaged values across blocks which may have obscured block-by-block effects.

Interestingly, rats showed reduced accuracy following the first reversal under all reward schedules. Furthermore, while rats performed less accurately in sessions with more uncertainty, behavioral changepoint and the number of perseverative errors following reversal did not vary with reward uncertainty or block. Experience with reversals is known to affect strategy and expectation of reversals (Costa et al., 2015; Mackintosh et al., 1968; Murray & Gaffan, 2006; Rudebeck et al., 2013; Yu & Dayan, 2005). We found that learning rate increased with experience over the first three blocks in the probabilistic schedules, indicating that the rats learned faster with repeated reversals. Conversely, inverse temperature, or choice determinism, decreased dramatically with increasing uncertainty but did not change in response to reversal experience. Inverse temperature affects stochasticity or the ratio of exploitation to exploration (Doya, 2002; Katahira, 2015), and the decrease in beta with uncertainty could be due to probability devaluation (Daw et al., 2006) or to increased exploration (Knox et al., 2012; Speekenbrink & Konstantinidis, 2015)

We also found that Win-Stay likelihood (sensitivity to positive feedback) was more strongly correlated with inverse temperature (Cinotti et al., 2019), while LS likelihood (sensitivity to negative feedback) was more strongly correlated with learning rate. However, both the WS likelihood and learning rate increased over repeated reversals, while inverse temperature and LS remained stable. It is unclear what is responsible for this mismatch, and this result comes in contradiction to Noworyta-Sokolowska and colleagues, who found that sensitivity to positive feedback is associated with faster learning (Noworyta-Sokolowska et al., 2019).

The second aim of this study was to determine whether the mOFC was critical for success in a blocked two-armed bandit task. The mOFC region that we targeted was also investigated by our lab in a study of progressive ratio responding (Swanson et al., 2019) and may be homologous to the pregenual anterior cingulate cortex in primates (Laubach et al., 2018). In the present study, we infused muscimol into mOFC to transiently inactivate the region. Inactivation led to a decrease in accuracy and a dramatic decrease in LS likelihood, as well as an increase in changepoint and perseveration. These results demonstrate that the mOFC plays an important role in TAB performance.

Previous studies showed similar reductions in negative feedback sensitivity following mOFC inactivation in a 80/20 performance-based two-armed bandit (Dalton et al., 2016; Verharen et al., 2020), though Dalton and colleagues also found a decrease in positive feedback sensitivity following mOFC inactivation. However, our results run contrary to a previous finding that mOFC inactivation decreases perseveration and increases Lose-Shift likelihood in a visual deterministic TAB task which used a 24/30 correct sliding window to trigger reversals (Hervig et al., 2020). This could be due to the difference in modality, as there is evidence to suggest that mOFC is more important in retrieving action-outcome associations than stimulus-outcome associations as compared to lateral OFC (Bradfield et al., 2015), and the animals in this experiment could be using a left-right strategy to solve the spatial bandit.

The final aim of this study was to explore the role of the noradrenaline system in a two-armed bandit task via the α2-noradrenoreceptor antagonist yohimbine. At lower doses than used here, systemic delivery of yohimbine has been shown to deactivate presynaptic α2-noradrenoreceptors, enhancing central NA release (Abercrombie et al., 1988; Szemeredi et al., 1991). For this reason, we also delivered yohimbine intracranially to determine the role of α2-adrenoreceptors in mOFC specifically (Agster et al., 2013; U’Prichard et al., 1979). It is worth noting that yohimbine antagonizes other monoamines including serotonin, (Millan et al., 2000), the signaling of which appears to play a general role in bandit performance (Izquierdo et al., 2017). Across our studies, we found similar effects of 2 mg/kg systemic yohimbine and 5 µg/µL intra-cortical yohimbine. Specifically, accuracy was reduced while changepoint increased under yohimbine in deterministic and 80%/20% probabilistic sessions.

Critically, these results demonstrate a dissociation between the effects of yohimbine NA blockade and broad inactivation of the mOFC. While both manipulations led to a decrease in accuracy, muscimol inactivation of mOFC led to a decrease in sensitivity to negative feedback via a selective decrease in LS likelihood and α2-noradrenoreceptor blockade in mOFC led to a decrease in positive feedback sensitivity. In a recent review, Wilson and colleagues proposed that the overarching role of orbitofrontal cortex is to represent task states (Wilson et al., 2014), leading to the common finding that lesion or inactivation leads to deficits in reversal learning but not initial discrimination learning (Schoenbaum et al., 2002; Rudebeck & Murray, 2008), as we found here.

The noradrenergic system is one of many neuromodulators of flexibility (Robbins et al., 2010; Robbins & Roberts, 2007) and is thought to play a role in mediating the explore/exploit tradeoff (Aston-Jones & Cohen, 2005; Daw & Doya, 2006; Doya, 2002; Yu & Dayan, 2005). Specifically, endogenous tonic NA rises following reversal (Aston-Jones et al., 1997), which is thought to facilitate exploration (Aston-Jones et al., 1999; Jepma & Nieuwenhuis, 2011; Shea-Brown et al., 2008) and signal to the OFC to update task state or context (Bouret & Sara, 2005; Dayan & Yu, 2006; Sadacca et al., 2017). In the present study, we found that rats performed better on blocks where the high-value option matched their initial choice (Figure 6). We also found a decrease in inverse temperature, a measure of the exploration/exploitation tradeoff. Yohimbine antagonism may lead to a decrease in tonic NA in mOFC which would block updating of context and therefore decrease responding to the non-preferred side.

Finally, the increased noradrenergic tone caused by systemic yohimbine can lead to increased measures of impulsivity in a variety of tasks (Mahoney et al., 2016; Sun et al., 2010; Swann et al., 2005). While we found an increase in speed and decrease in accuracy under yohimbine, these measures varied independently based on correlation analysis.

We acknowledge several limitations and alternative interpretations for this study. First, we used only male rats in this study. We originally included both males and females, but the females performed fewer trials and did not reach the accuracy thresholds. However, this is inconsistent with published research and there is evidence for sex differences in strategy in probabilistic bandits (Chen et al., 2021). It is possible that these results do not generalize to both sexes.

Second, correlated reward schedules were presented in order of certainty and each schedule was presented one session per day for three days in a row. While we found no difference in strategy by day for each schedule, as reported by Noworyta-Sokolowska et al. (2019), it is possible that the order of presentation had some effect on WSLS strategy. Also, we only report the first 6 blocks in all block-based analyses although some rats completed up to 15 blocks per session. Focusing on the initial part of the test session avoided potential effects of satiety (Colwill & Rescorla, 1985; Rudebeck & Murray, 2011). We note that performance remained relatively stable until the rats approached the end of the one-hour session and reached satiety. We further note that we removed any incomplete blocks in the whole-session analysis.

Third, rats were tested only in the deterministic reward schedule to increase the likelihood of detecting an effect. We were therefore unable to determine if the effects muscimol inactivation of OFC depended on reward uncertainty. Previous studies that implemented a performance-based bandit used an 80/20 probabilistic schedule (Dalton et al., 2016; Verharen et al., 2020) and we found that baseline LS likelihood in the blocked bandit depends on the presence of uncertainty. There is also some evidence that the result depends on species, since excitotoxic, fiber-sparing lesions of OFC in monkeys do not impair deterministic reversal learning (Rudebeck et al., 2013).

Fourth, we found a contradiction by scope of the RL model. When looking at each block individually, systemic yohimbine decreased learning rate but had no effect on inverse temperature. Inverse temperature controls the influence of reward history (Katahira, 2015), so the difference in evidence available for a single block and for the whole session likely contributed to the difference in fits for alpha and beta. Within such a short window, a decrease in likelihood of selecting the higher-value option could just as well be attributed to slow updating of the value as to the reduced influence of the value

Finally, the side-biased Win-Stay likelihood and accuracy under yohimbine in the deterministic session could be due to a failure in spatial encoding. As discussed above, yohimbine can increase central NA release (Abercrombie et al., 1988; Szemeredi et al., 1991), so the dose used in this study may increase activation of β-noradrenoreceptors, which control synaptic plasticity in hippocampus (Hagena et al., 2016; Kemp & Manahan-Vaughan, 2008). In addition to reduced activity in the frontal cortex (Kovács & Hernádi, 2003; Zhang et al., 2013) systemic yohimbine may lead to altered spatial representations in the hippocampal formation (Grella et al., 2019; Wagatsuma et al., 2018). Yohimbine may therefore impair the ability to form new spatial/action-outcome contingencies. However, the intra-cortical infusions also resulted in this pattern of reduced performance in alternate blocks. Since the infusion would not have affected LC somatic autoreceptors, this would not lead to an increase in central NA as could be caused by systemic yohimbine (Huang et al., 2012). However, this pattern did not appear in the 80/20 session and since the cannulated animals weren’t tested under this schedule, it is difficult to confirm how uncertainty relates to the role of noradrenaline in context updating.

## Notes

**Author Note** We have no conflicts to disclose. This work was supported by a Faculty Research Support Grant and Doctoral Dissertation Award from American University to ML and KS, respectively. We thank Drs Vincent Costa, David Kearns, Elizabeth Murray, and Catherine Stoodley for helpful comments on this study and manuscript.

### Competing Interest Statement

The authors have declared no competing interest.

### Summary of Updates

Revisions were made to address reviewer comments from reviews at the Journal of Neuroscience and Behavioral Neuroscience. A major change in the manuscript was more stringent statistical analyses, which revealed effects of systemic but not intra-cortical yohimbine on behavior in a two-armed bandit task.

